# Towards measuring growth rates of pathogens during infections by D_2_O-labeling lipidomics

**DOI:** 10.1101/330464

**Authors:** Cajetan Neubauer, Alex L. Sessions, Ian R. Booth, Benjamin P. Bowen, Sebastian H. Kopf, Dianne K. Newman, Nathan F. Dalleska

## Abstract

**RATIONALE:** Microbial growth rate is an important physiological parameter that is challenging to measure *in situ*, partly because microbes grow slowly in many environments. Recently, it has been demonstrated that generation times of *S. aureus* in cystic fibrosis (CF) infections can be determined by D_2_O-labeling of actively synthesized fatty acids. To improve species specificity and allow growth rate monitoring for a greater range of pathogens during the treatment of infections, it is desirable to accurately quantify trace incorporation of deuterium into phospholipids.

**METHODS:** Lipid extracts of D_2_O-treated *E. coli* cultures were measured on LC-ESI-MS instruments equipped with TOF and Orbitrap mass analyzers, and used for comparison with the analysis of fatty acids by isotope-ratio GC-MS. We then develop an approach to enable tracking of lipid labeling, by following the transition from stationary into exponential growth in pure cultures. Lastly, we apply D_2_O-labeling lipidomics to clinical samples from CF patients with chronic lung infections.

**RESULTS:** Lipidomics facilitates deuterium quantification in lipids at levels that are useful for many labeling applications (>0.03 at% D). In the *E. coli* cultures, labeling dynamics of phospholipids depend largely on their acyl chains and between phospholipids we notice differences that are not obvious from absolute concentrations alone. For example, cyclopropyl-containing lipids reflect the regulation of cyclopropane fatty acid synthase, which is predominantly expressed at the beginning of stationary phase. The deuterium incorporation into a lipid that is specific for *S. aureus* in CF sputum, indicates an average generation time of the pathogen on the order of one cell doubling per day.

**CONCLUSIONS:** This study demonstrates how trace level measurement of stable isotopes in intact lipids can be used to quantify lipid metabolism in pure cultures and provides guidelines that enable growth rate measurements in microbiome samples after incubation with a low percentage of D_2_O.

## INTRODUCTION

Bacteria continually react to diverse stimuli, such as the availability of nutrients and electron acceptors, exposure to antimicrobial drugs or attack by the immune system. However, measuring microbial metabolites and growth rates within a complex environment still poses many technical challenges. Two recent advances in microbial ecology are beginning to make measuring average growth rates in environmental samples possible. The first advance is based on metagenomic DNA sequencing and takes advantage of the observation that growing cells yield more sequencing reads at genomic regions near the origin of replication ^[1,2]^. This method is applicable to any microbial species in a microbiome as long as its assembled genome has a high sequence coverage. The second advance uses isotopic labeling to determine the biosynthesis rates of microbial lipid metabolites by mass spectrometry ^[3,4]^. Stable-isotope probing has a larger dynamic range than sequencing and can be used to quantify the slow growth rates that microbes have under environmental conditions. A limitation of stable-isotope probing, however, is the identification of metabolites that are diagnostic for a specific microorganism. It is therefore desirable to combine isotopic labeling with a method such as lipidomics, which can detect a large number of microbial metabolites.

Lipids have been used for decades in ecology as markers of microbial metabolism, where they reveal information about viable biomass, nutritional status or changes of the microbial community structure ^[5,6]^. Also, lipids can still be analyzed long after nucleic acids and peptides are degraded ^[7]^. In order to estimate the growth rates of microbes, the active production of strain- and genus-specific lipid metabolites can be measured with stable isotope labeling ^[3]^. With advances in soft-ionization mass spectrometry we can now attempt to combine isotope quantification and lipidomics for the study of microbes *in situ*.

Soft-ionization mass spectrometry detects thousands of lipids in environmental extracts and would in principle be well-suited to quantify the biosynthesis of lipid biomarkers by itself ^[8–11]^. However, extraction yields and ionization efficiency vary widely between samples ^[12–16]^. This currently poses severe constraints on measuring lipid production rates and effectively limits estimating bacterial growth rates in natural environments.

Ratios of isotopes can be measured with high accuracy by mass spectrometry, partly because ratiometric readouts vary less than absolute ion intensities ^[17]^. For lipid biosynthesis, the incorporation of an isotope tracer such as ^13^C-labeled substrates hence can provide a robust way to quantify anabolic activity and lipid turnover ^[18,19]^. In microbiome samples bacteria differ widely in their ability to take up carbon sources and gases such as CO_2_, depending on their genetic capabilities and metabolic states. D_2_O is a non-discriminating tracer of *de novo* lipid biosynthesis and thus often better suited for microbiome studies. D_2_O-labeling has recently been used to estimate *in situ* growth rates of *Staphylococcus aureus* in chronically infected lungs. After labeling of expectorated sputum with D_2_O, the deuterium enrichment of *anteiso* fatty acids was quantified using GC pyrolysis isotope-ratio MS (GC/P/IRMS) in order to estimate the growth rate of the pathogen ^[3]^. Previous studies have been used to study lipid biosynthesis with D_2_O *in vivo* ^[20–25]^. Environmental samples, however, provide particular challenges. For example, microbes often grow slowly *in situ* and one can expect low rates of deuterium incorporation into lipids ^[3]^. Whether trace levels of incorporation can reliably be detected remains to be studied before deuterium incorporation can be used to determine lipid biosynthesis rates *in situ* by lipidomics.

In this study we apply stable isotope probing with D_2_O and measure deuterium incorporation by MS-based lipidomics. This approach can be used to obtain labeling rates for individual intact lipids in environmental samples, where microorganisms typically grow slowly. We begin by characterising several technical aspects that have to do with quantifying low levels of deuterium labeling. We then refine our application of D_2_O-labeling lipidomics by tracking of lipid labeling in *E. coli* during the stationary-to-log phase transition. This reveals lipids that have distinct labeling dynamics that are not obvious from measuring absolute analyte concentrations alone. Lastly, we test D_2_O-labeling lipidomics in a clinical context with the aim to measure the growth of *S. aureus* in cystic fibrosis lung infections. In sum this study establishes principles for how growth rates of microbes *in situ* can be estimated by stable isotope probing lipidomics.

## RESULTS

When cells grow after addition of heavy water, newly synthesized biomass will contain more D. This also means that each lipid pool will be a mixture of molecules that vary more in their D abundance. The introduced heterogeneity causes broadening of chromatographic peaks, which could skew the isotope ratio observed by LC-MS as ionization efficiency varies over time ^[26,27]^. In order to evaluate how this affects isotope quantification by lipidomics, we grew an *E. coli* culture in 4 % D_2_O (fractional D-abundance, ^2^*F*_WATER_) and measured lipids after chromatographic separation using an ESI-TOF mass spectrometer ^[28]^.

*E. coli* has a comparatively simple lipid composition and its lipid metabolism has been studied for decades ^[29]^. The bacterium therefore provides a solid model system to develop and test methods for stable isotope labeling lipidomics. *E. coli* lipid extracts contains mainly phosphatidylethanolamines (PE), phosphatidylglycerol (PG) and cardiolipins (CL) ^[30]^. When fully labeled in 4 % D_2_O, the molecular ions from PE and PG lipids extend over a range of 8 *m/z*. As expected, labeling causes a shift of retention time (Figure 1A). Strongly deuterated molecules elute earlier than lighter ones (Figure 1B). The maximum shift in retention time is about half of the chromatographic peak width, which indicates that all molecular species have overlapping elution profiles (Figure 1C). We expect that this degree of shifting typically does not alter isotope ratios, as long as a moderate amount of labeling is used and the mass spectrum is integrated over a sufficiently large retention time window.

**Figure 1:**
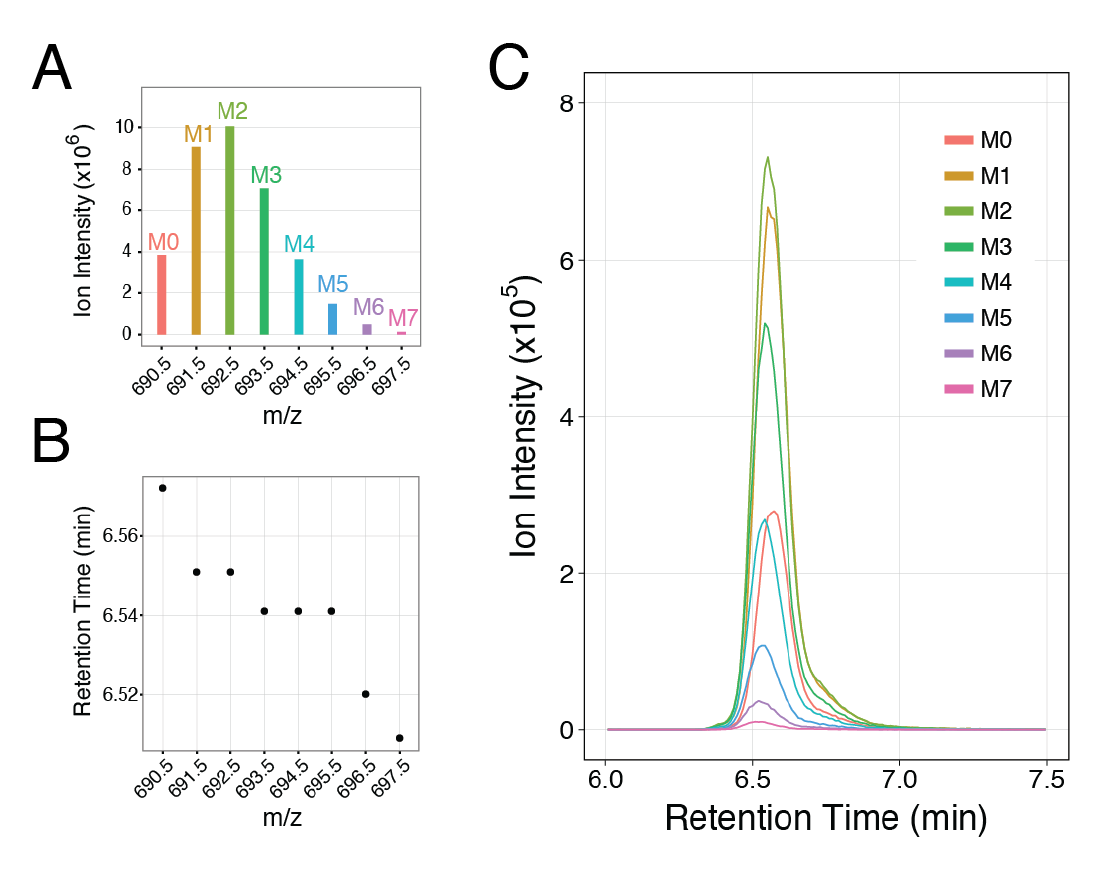
Deuterium-labeling shifts the chromatographic retention of lipids. LC-MS data of labeled PE(16:0/16:1), made by an *E. coli* culture grown in 4 % D_2_O. The retention time shift was evident, yet most isotopologues co-eluted and derived isotope-ratios were not altered. **(A)** Isotopologue distribution (M0 to M7; *m/z* ± 0.05). Data collected in positive mode. **(B)** Retention time at maximum ion intensity for each *m/z* range. Spectra were collected at a rate of 95 scans per minute. **(C)** Extracted ion chromatograms for the eight *m/z* ranges.

The quantification of D in intact lipids is complicated by ^13^C, which is naturally present in lipids at about 1.1 %. Mass gained by ^13^C or D cannot be distinguished by current TOF mass analyzers (resolving power ~30,000). Resolving the minute mass difference (~3 mDa) is possible for small lipids (<600 Da) by Orbitrap MS, but this approach is currently not ideal for LC-MS due to the long scan times (~1 second). So we need a procedure to determine the gain of isotopic label indirectly by comparing lipid extracts from bacterial cultures grown with and without label. The average molecular weights of the two mass distributions are calculated and their difference, *ΔMW*, interpreted as the mass gained by D incorporation (Figure 2). The fractional abundance of D in a lipid (^2^*F*_LIPID_) is then calculated by dividing *ΔMW* by the number of C-bound hydrogens. Using this method, glycerophospholipids produced by *E. coli* grown in 4 % D_2_O yield a ^2^*F*_LIPID_ of about 2.5 %. Values lower than 4 % are expected because hydrogen atoms from the unlabeled carbon source are incorporated into the lipids and biosynthetic enzymes favor ^1^H over D due to kinetic isotope effects. Note that this calculation assumes that N- and O-bound hydrogen atoms equilibrate fully with water during extraction and chromatography ^[31]^. Additionally, the natural level of D, which is about 0.015 %, is neglected for the purpose of this study.

**Figure 2:**
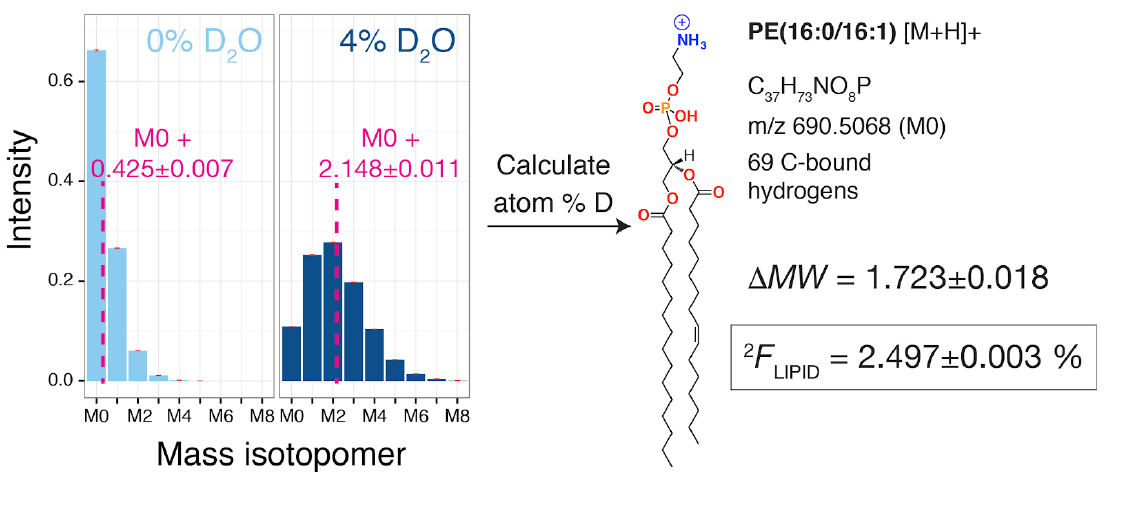
Calculating the deuterium content of intact lipids from an unlabeled and labeled sample. Two cultures of *E. coli* FRAG-1 were grown without and with D_2_O addition (4 %). The molecular weight was calculated from each distribution and is shown relative to M0 in magenta (±1σ). The deuterium content of the lipid was calculated by dividing the mass difference *ΔMW* by the number of C-bound hydrogens (for details see Methods). ^2^*F*_LIPID_ generally had a standard deviation < 0.005 at%. The isotopic distribution of PE(16:0/16:1) in positive mode is shown. Lipid extracts measured by LC-MS using injection volumes of 1, 3 and 9 μL. Note that error bars (±1σ; magenta) are small.

In order to evaluate the utility of D_2_O-labeling lipidomics for estimating bacterial growth rates, we grew *E. coli* cultures in glucose minimal medium ranging from 0.0156 % (natural abundance) to 4 % ^2^*F*_WATER_ and quantified the glycerophospholipids PE and PG. ^2^*F*_LIPID_ increases linearly (R^2^ > 0.99) with ^2^*F*_WATER_ (Figure 3A and B). Analyzing the same samples on a Q Exactive Plus Orbitrap operated at R=35,000, yields nearly identical slopes. Assuming a detection limit of 0.03 % ^2^*F*_LIPID_ for D_2_O-labeling lipidomics, we suggest that incubating cells for 15 minutes with 5 % ^2^*F*_WATER_ is a useful range to quantify lipid biosynthesis from microbes growing at one doubling per day (Figure 3C). These boundary conditions indicate that D_2_O-labeling lipidomics can be developed further into a method to estimate microbial growth rates in environmental samples ^[3]^. An important consideration for microcosm incubations is that two separate populations of molecules co-occur after labeling, a pool that contains low natural D abundance and a new pool that is enriched in D. High labeling strength would create molecules that occur further away from the monoisotopic mass in the spectrum and become difficult to quantify.

**Figure 3:**
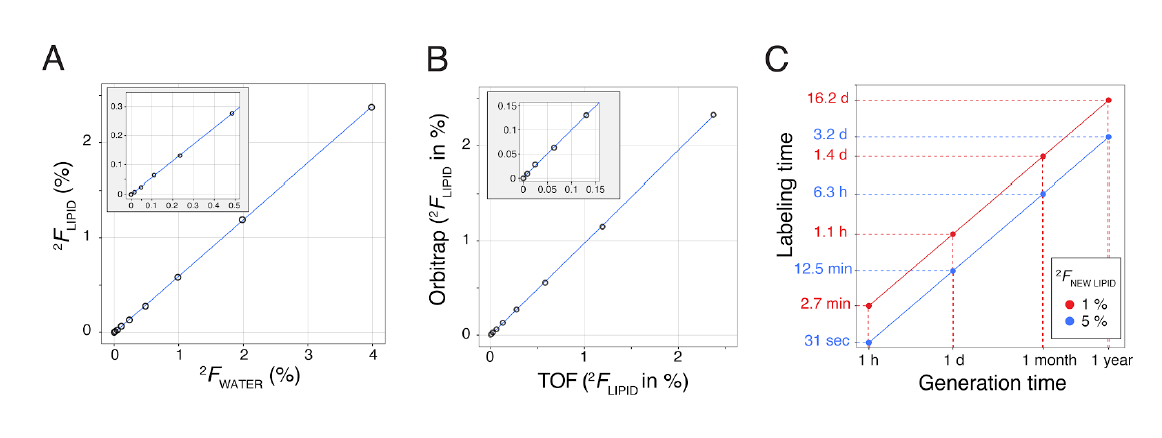
Isotope-ratios by lipidomics are suitable for measuring microbial growth rates *in situ*. (A) Overnight cultures of *E. coli* were grown in M9-glucose medium containing varying amounts of D_2_O and lipid extracts analyzed by LC-MS. The measured deuterium content of PE(16:0/16:0(Cp)) is shown. Enrichments with ^2^*F*_LIPID_ > 0.03 % were within the linear range. **(B)** Comparison of a Xevo G2 TOF instrument and a Orbitrap Q Exactive Plus (resolving power 35,000 at m/z 200) using the same samples as in (A). ^2^*F*_LIPID_ values for PE(16:0/16:0(Cp)) are plotted. The linear regression has a slope of 0.976 and R^2^ of 0.999. **(C)** Estimate of how long D_2_O-labeling has to be performed to achieve deuterium enrichment of +0.03 at% D in lipids, assuming that the newly made lipid fraction (^2^*F*_NEW LIPID_) gets labeled at 1 % (red) or 5 % (blue). For a microbe growing with a doubling time of one day, incubation would need to be performed for 12.5 min or 1.1 hours, respectively, to reach en enrichment of 0.03 % over natural ^2^*F*_LIPID_. We here estimate the enrichment at time t using the following equation: ^2^*F*_LIPID_ (t) = ^2^*F*_NEW LIPID_ * (1 2^t/GT^).

Ionization conditions can affect isotopologue distributions and thus alter isotope ratios ^[32]^. We varied the injected sample amount, ionization mode, capillary voltage, desolvation temperature and desolvation gas flow without noticing significant changes. For instance, PE(16:0/16:1) had a ^2^*F*_LIPID_ of 2.549±0.003 (1σ) in positive ionization mode and 2.567±0.019 in negative mode (lower precision due to increased background noise). Less abundant analytes have greater standard deviations. PE(16:0/16:0), which was 10-times less abundant, had a ^2^*F*_LIPID_ of 2.432±0.078 (positive mode). These trials show that the D abundance of lipids can be measured reproducibly. However, the most intense signals must be within the linear range of the mass analyzer and the detector must not be in dead time on a TOF instrument. Also, isotopologue patterns that are affected by co-eluting compounds have to be excluded. In our UPLC setup this was the case for some cardiolipins (m/z >1,200).

For the calculation of ^2^*F*_LIPID_ we assume that C-bound hydrogens do not exchange with solvent water during sample preparation and electrospray ionization, while N- and O-bound hydrogens fully equilibrate. If this is not the case, we would obtain inaccurate ^2^*F*_LIPID_ values ^[33,34]^. To test for H/D exchange we compare UPLC-ESI-TOF with GC/P/IRMS, which quantifies near-natural isotopic composition of fatty acids ^[35]^. Albeit the two methods are distinct in many ways, they should yield a similar linear relationship between ^2^*F*_WATER_ and ^2^*F*_LIPID_ ^[36,37]^. For lipidomics, we determine an average slope for intact lipids produced by *E. coli* in glucose minimal medium of 0.577±0.003 (Figure S1; ^2^*F*_WATER_ between 0.125 and 4 %). Slightly higher slopes have been reported previously for *E. coli* fatty acids using GC/P/IRMS (0.65±0.04 for C16:0, 0.60±0.02 for C16:1 and 0.63±0.03 for C18:1) ^[37]^. Growth on acetate raises ^2^*F* in *E. coli* fatty acids analyzed by GC/P/IRMS, and it does so also for intact lipids measured by LC-MS (Figure S1). Overall, we obtain comparable slopes by lipidomics and GC/P/IRMS.

In order to further constrain H/D exchange, we dissolved PE(18:0/18:1) in acetonitrile, added H_2_ O or D_2_O (sold as 99.9 at% D), and recorded mass spectra by direct infusion. The addition of D_2_O shifts the mass spectrum by four units in positive ionization mode, as expected for an analyte that has four non C-bound hydrogens (Figure 4). The *ΔMW* of 3.86 suggests that the four exchangeable hydrogens in PE(18:0/18:1) [M+H]+ have an average probability of 97.4 % to contain D. A theoretical spectrum that assumes four positions in the ion to have this probability for D closely matches the measured spectrum. Importantly, D_2_O addition does not yield detectable signal beyond a shift of the unlabeled distribution by four mass units. Such isotopologues would occur if the exchange of C-bound hydrogens occurs at high rates during ESI. Absence of these signals indicates that C-bound hydrogens exchanged at least 2000-fold slower than non-C-bound hydrogens, which is in line with prior assessments of C-bound hydrogen exchange ^[38]^. Together these tests imply that for lipids labeled well above natural D-abundance, no relevant artifacts of ^2^*F*_LIPID_ values due to exchange of C-bound hydrogen are likely in UPLC-ESI-TOF.

**Figure 4:**
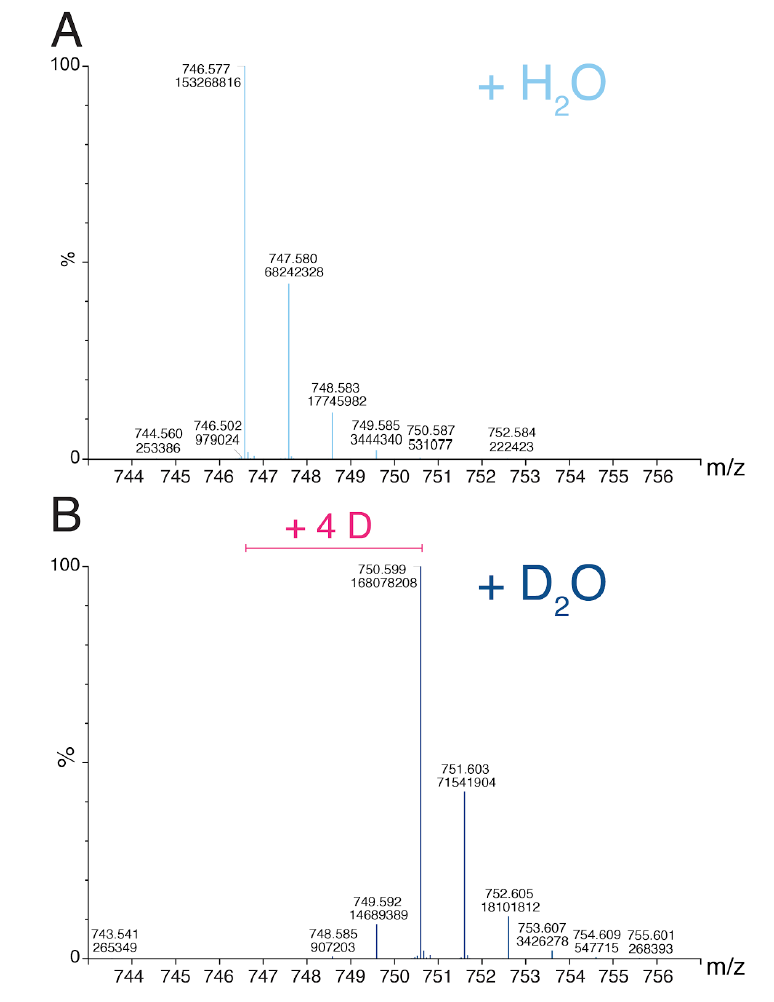
Direct infusion of lipid in the presence of H_2_ O or D_2_O showed no exchange of C-bound hydrogens. Direct infusion mass spectra of PE(18:0/18:1) ([M+H]+ ion) dissolved in **(A)** H_2_ O (0.015 at% D) or **(B)** D_2_O (99.9 at% D). The molecular weight shift *ΔMW* was 3.86 and indicated that maximally 97.4% of the four exchangeable sites exchanged to deuterium. No signal above background was detected beyond *m/z* 750.6 (H_2_ O sample) and *m/z* 754.6 (D_2_O sample). The solution contained a 4:1 ratio of acetonitrile to water and was buffered with 10 mM ammonium formate and 0.1% formic acid, which contained ^1^H. Data was acquired for 4 minutes at 0.4 mL/min flow rate.

With the addition of small quantities of D_2_O to pure cultures we have an opportunity to measure lipid isotope ratios and absolute concentrations simultaneously and compare the two quantifications side by side. Our test case here is the lipid metabolism of *E. coli* during the transition from stationary phase into exponential growth. Cells from two stationary phase precultures (*u*:‘unlabeled’ and *l:*‘labeled’ in 4 % ^2^*F*_WATER_) were used to inoculate two cultures each of unlabeled (*U*) or labeled (*L*) medium (Figure 5). Four growth cultures (*uU, lU, uL, lL*) were sampled to determine optical density (OD_600_), as well as bulk protein and lipid concentrations. During the 160 minutes of labeling, cells divided three to four times (Figure S2).

**Figure 5:**
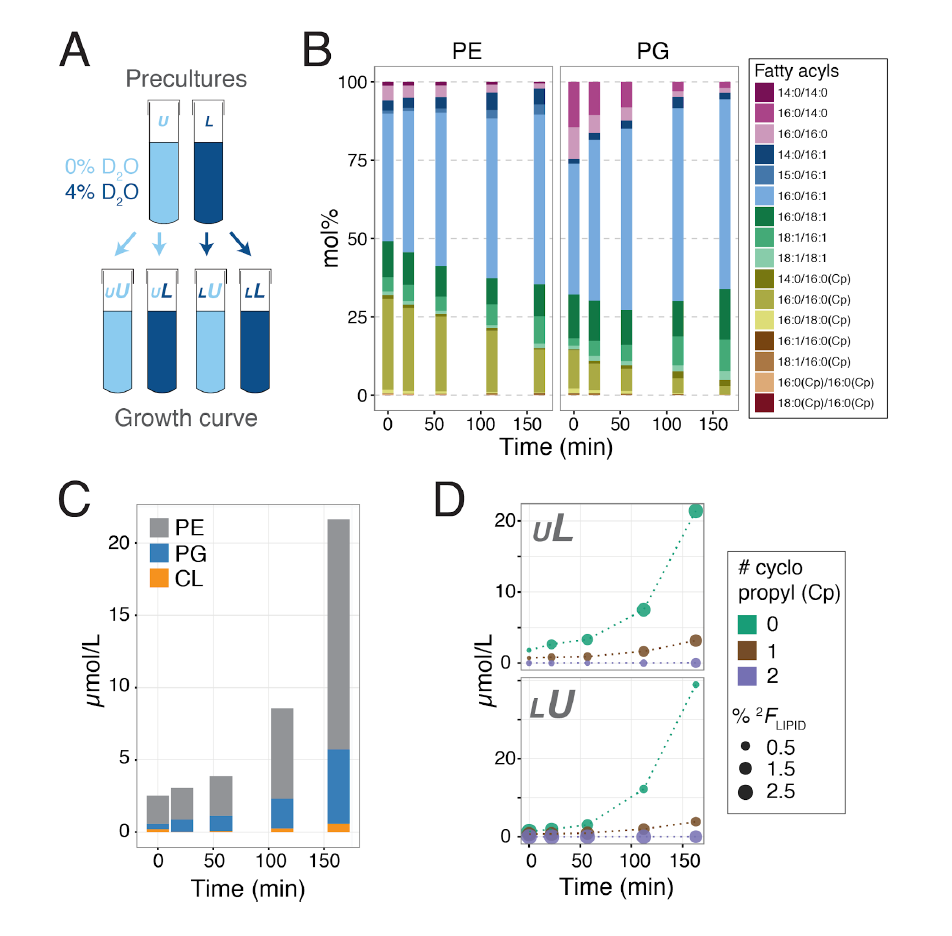
Lipid production in *E. coli* during stationary phase exit was followed by absolute concentrations and deuterium incorporation. (A) Four cultures were followed by lipidomics. Labeling scheme: *l* = labeled (4 % D_2_O; dark blue) stationary preculture, *u* = unlabeled (light blue) stationary preculture, *L* = labeled (4 % D_2_O) growth culture, *U* = unlabeled growth culture. **(B)** Relative abundance of PE and PG lipids. Cyclopropane fatty acyls are more abundant in stationary phase compared to growth phase (data from *uL* culture). (C), double bonds, and cyclopropane rings (Cp) in acyl chains. **(C)** Quantification of lipid classes (data from *uU* culture). Throughout the outgrowth the proportion of PE (grey) was 70-78 mol% and PG (blue) 19-24 mol%. CL (orange) is synthesized from two molecules of PG and often increased in stationary phase ^[56]^. In this time course CL was 3.5 mol% in the inoculum and 1.5-2.5 mol% during outgrowth. **(D)** Time course for PE and PG lipids with 0 (green), 1 (brown) or 2 cyclopropyl rings (violet) in cultures *uL* and *lU*. Size of the data points represents deuterium abundance in the lipid (^2^*F*_LIPID_).

In these tests, the stationary phase *E. coli* cultures contain a high proportion of cyclopropane fatty acids (CFA). Greater than 25 mol% of PE and PG phospholipids contain at least one acyl chain with cyclopropyl ring. When cells resume growth, CFA abundance decreases to about 12 mol% (Figure 5B and C). The formation of CFAs in *E. coli* is a post-synthetic modification of the unsaturated phospholipids that occurs predominantly as cultures enter the stationary phase. CFA synthase has an unusual regulation that involves enzyme instability as well as transcription of the *cfa* gene from two distinct promoters ^[39,40]^. This means that, although CFA synthase is synthesized at basal levels throughout the growth curve, a transient spike in activity occurs during the log-to-stationary phase transition. In agreement with this regulation, CFAs largely dilute out during stationary-to-log phase transition. Using D_2_O-labeling lipidomics we detect small levels of production of CFA lipids as well as D incorporation, which shows that CFA lipids were actively made during stationary phase exit (Figure 5D).

Untargeted labeling reveals striking differences between phospholipids. Here we describe the *uL* scenario in detail. Some D_2_O-labeling patterns fit an exponential growth model (Figure 6). Other lipids, in particular CFA-containing lipids, were inconsistent with simple exponential *de novo* production. For them the growth model needs to be extended. We include a parameter that accounts for lipid biomass in the inoculum that is inactive, i.e. not exponentially reproduced during the stationary-to-log phase transition (see Methods for details).

**Figure 6:**
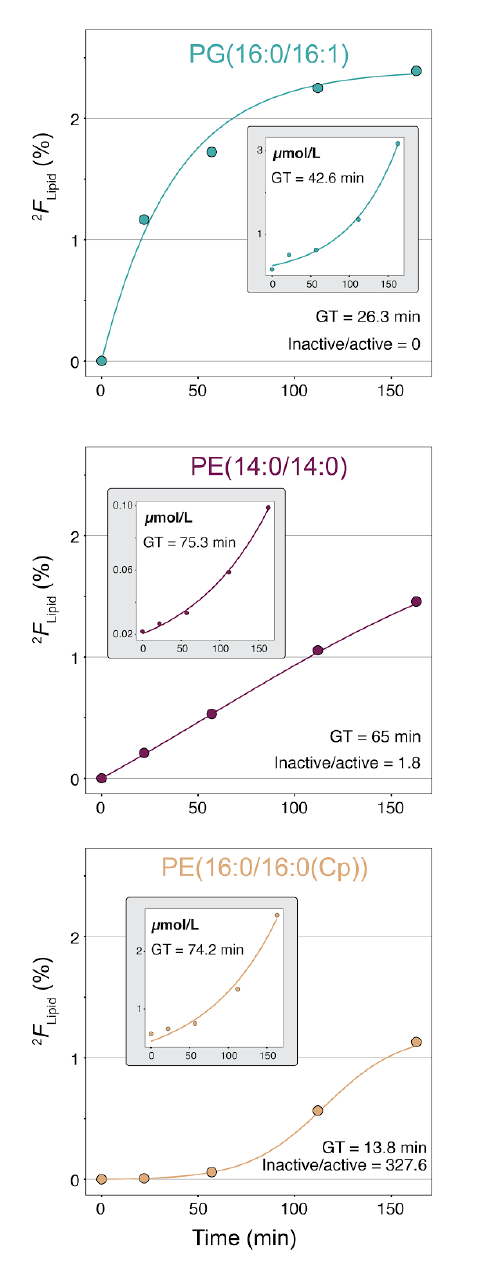
During stationary phase exit lipids have distinct labeling dynamics, which are dominated by the fatty acyl chains. The deuterium abundance (^2^*F*_LIPID_) in three representative lipids is shown (data from culture *UL*). Lines represent the best fit of a growth model that estimates generation time (GT) and the ratio of inactive to active unlabeled material at time 0 (see Methods for details). The insets show absolute concentrations for comparison. See Figure S3 & S4 for a comprehensive overview of lipids.

The isotopic labeling patterns of *E. coli* phospholipids are dominated by their two fatty acyl chains, as they contain most of the C-bound hydrogens. A major trend we notice is that lipids that contain unsaturated fatty acids label rapidly, while fully saturated lipids incorporate label more steadily (Figure 6; also Figure S3 and S4). In *E. coli*, unsaturated fatty acids are made during *de novo* fatty acid biosynthesis and not generated by modification of saturated fatty acids or phospholipids ^[29]^. The faster labeling of unsaturated lipids we observe thus likely reflects that the unlabeled inoculum contained little unsaturated phospholipids, because most got converted into CFA during stationary phase. A second common trend is that most CFA-containing lipids show slow initial increase of ^2^*F*_LIPID_ and often do not reach full saturation levels. This reflects that CFA lipids are only produced in small quantities after inoculation and hence a large proportion of unlabeled material is carried over from stationary phase. As CFA formation is a post-synthetic modification, labeling of CFA lipids additionally depends on the prior labeling of the precursor pool.

Interestingly, the two common trends we observe, namely slower labeling of saturated lipids compared to unsaturated lipids and slow and incomplete labeling of CFA-containing lipids, do not apply to all phospholipids. For example, PE(16:0/18:1) and PG(16:0/18:1) have distinct labeling patterns (Figure S3). Generally, the labeling of PE lipids that have one unsaturated and one saturated straight chain fatty acyl reveal a significantly larger proportion of unlabeled lipid compared to their PG analogs. Distinct labeling dynamics also occur for some CFA lipids. While most CFA lipid pools label slowly and do not reach high labeling, production of PG(14:0/16:0(Cp)) is stimulated so that it gains label rapidly and to high levels (Figure 6). This lipid occurs only in trace amounts in stationary phase, as does its precursor PG(14:0/16:1). Therefore, the material produced during outgrowth of the cultures is highly labeled and dominates the PG(14:0/16:0(Cp)) pool. Overall, these tests indicates that D_2_O addition allows a readout of how much of the material has been newly synthesized even for minority components, whose absolute concentrations can be challenging to quantify in complex lipid extracts.

The results so far indicate that lipidomics can be used to measure bacterial lipid biosynthesis in pure cultures. If D_2_O-labeling lipidomics could quantify microbial growth reliably *in situ*, this might for example enable the use of microcosm incubations to test how different drugs impact the microbial community of individual patients. A disease context that is well-suited to assess the practicability of D_2_O-labeling lipidomics for complex samples are cystic fibrosis (CF). These chronic lung infections contain heterogeneous mixtures of human biomass and microorganisms. Some of the bacteria in CF lungs become pathogenic and tend to develop drug-resistant phenotypes. In previous work on D_2_O-labeled expectorated CF sputum we have examined the growth of *S. aureus* via D/H ratios of *anteiso*-fatty acids, an abundant fatty acids of this pathogen (3). It is important, however, that *anteiso*-fatty acids are produced also by other bacterial species. In the context of CF sputum, *Prevotella melaninogenica* and *Stenotrophomonas maltophilia* are relevant sources of *anteiso-*C15:0 and *anteiso-*C17:0 fatty acids in some CF patients ^[3,41,42]^. Certain phospholipids, specifically those that contain *anteiso* fatty acyls, may therefore be more specific markers of *S. aureus* in CF infections and could be used to assess activity of the pathogen by lipidomics. To evaluate this hypothesis, we analyzed samples that had been collected and characterised as part of a longitudinal study of CF patients undergoing pulmonary exacerbations ^[42]^.

A lipid that is appears well-suited to monitor the growth of *S. aureus* is PG(*a-*C15:0/*a-*C17:0). This compound was detected in lipid extracts of *S. aureus* and its structure assigned based on the *m/z* of the molecular ion in positive and negative ionization mode as well as MS/MS fragmentation spectra. Subsequently, signals from this lipid were also detected in expectorated sputum from several CF patients with *S. aureus* infection (Figure 7). Two patient whose lung infections did not contain *S. aureus* showed no signal corresponding to PG(*a*-C15:0/*a-*C17:0). Based on these observations PG(*a-*C15:0/*a-*C17:0) in CF sputum appears to be an specific marker for *S. aureus* in CF sputum.

**Figure 7:**
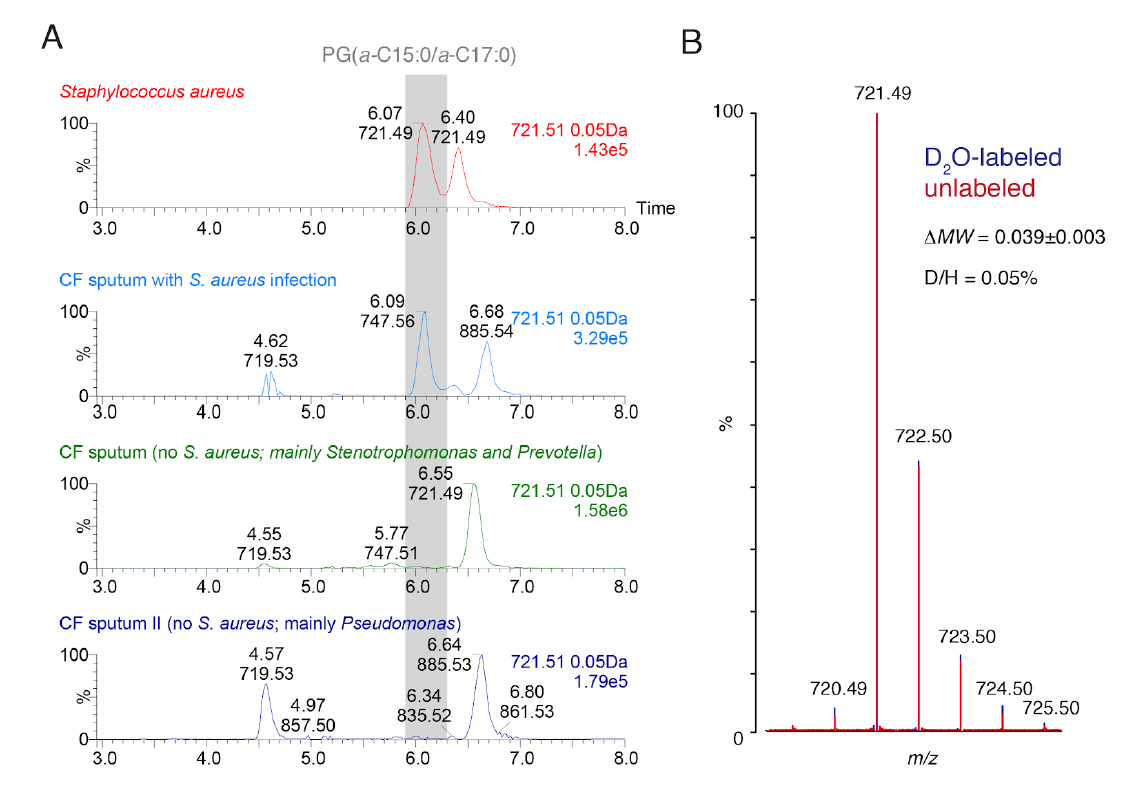
Growth of *S. aureus* in cystic fibrosis (CF) lung infections. (A) *S. aureus* (top) produces phospholipids that contain *anteiso* fatty acids. PG(*a-*C15:0/*a-*C17:0), which has a *m/z* 721.51 in negative ionization mode and elutes at 6.1 min, is shown as example. In expectorated sputum from a CF patient with a *S. aureus* infection (second from top) PG(*a-*C15:0/*a-*C17:0) is detected, while no comparable levels are detected in a controls from two patients whose infection does not contain *S. aureus* (bottom two panels). Note that *S. aureus* analytes occur only at trace abundances in sputum and concentrated lipid extracts were used to collect mass spectra. **(B)** Deuterium content of PG(*a-*C15:0/*a-*C17:0) from CF sputum labeled for 1 hour at 4 % D_2_O (blue) and unlabeled sputum (red). From the D/H of 0.05% generation times of *S. aureus* on the order of one cell doubling per day can be inferred ^[3]^.

Microbial lipid metabolites make up only a minute fraction of the total lipid content of CF sputum. The high sensitivity of ESI mass spectrometry allows detection of trace components, however, we had to use concentrated lipid extracts to yield sufficiently high signal intensities for the target analyte. In order to minimize contamination of the mass spectrometer, we only collected MS data at retention times that are needed to detect abundant phospholipids of *S. aureus* (5-8 min.). Labeling of CF sputum with 4% D_2_O for 1 hour resulted in 0.054±0.04 at% D enrichment of PG(*a-*C15:0/*a-*C17:0). This value can be used to estimate that the average generation time of *S. aureus* was approximately one cell doubling per day. This estimation is based on a previously established procedure that takes into account diffusion of the label, cell maintenance and other factors ^[3]^. For comparison, the generation time estimated by the D/H ratio of *anteiso-*C15:0 fatty acid in this sample was 3.3 days ^[42]^. The slower estimate based on GC isotope-ratio MS could for example be caused by contributions of *anteiso-*C15:0 from other sources or variability in the production rates of *anteiso*-C15:0 containing phospholipids in *S. aureus*. In summary, these initial tests indicate that it is possible to measure the activity of microbial pathogens *in situ* by D_2_O-labeling lipidomics. Its main benefits are that LC-MS has increased species specificity, requires smaller sample amounts, it is faster than alternative MS methods ^[43]^. Furthermore, D_2_O-lipidomics can be performed on instrumentation that is available in many biomedical laboratories.

## CONCLUSIONS

The combination of D_2_O-labeling and lipidomics allows a robust isotope ratio measurement, which reveals dynamic aspects of biosynthesis not accessible from absolute concentrations alone. The technology is also sufficiently sensitive to be adapted for environmental samples. Based on this study we encourage the development of LC-MS assays for the analysis of microbial growth in microbiome samples. Routine methods to measure bacterial growth in clinical samples are important to better understand microbial physiology in infections and improve diagnostics.

A critical parameter for D_2_O-labeling lipidomics is labeling strength. Very high concentrations of D_2_O (e.g., 20 to ~100 %) are tolerated by microorganisms and can be used to monitor biosynthesis ^[3,4]^. For LC-MS high labeling strengths are not desirable because they cause broad isotopic distributions. Quantification would become especially difficult when only a small proportion of the lipid is newly produced. In such a scenario, the labeled lipid would have a broad mass distribution far from the monoisotopic mass and potentially even overlap with other compounds. Interestingly, the ratio M1/M0 increases approximately linearly with ^2^*F*_LIPID_ (Figure S5). M1/M0 could be a simple readout of D incorporation in environmental samples. When we assume an excess of unlabeled over labeled lipid, as it is the case for many environmental incubations, we anticipate an optimal labeling strength that causes the greatest change of the M1/M0 ratio. This is achieved when the *ΔMW* of the newly-made lipid is about +1.5 Da. Overall, a concentration of 2-3 % ^2^*F*_WATER_ seems most suited for environmental microcosm incubations. The optimal value will depend on the complexity of the lipid sample, i.e. whether D incorporation can be assessed from isotopologue distributions or M1/M0 ratio. Another consideration is that the fraction of D that enters the lipid varies with microbial metabolism ^[37,44]^. We estimate that the combination of D_2_O-labeling and lipidomics as used here can roughly quantify growth rates greater than one cell doubling per day after labeling for 15 minutes.

D_2_O-lipidomics can in principle track lipid biosynthesis for many lipids in the same way as we have done here for 27 abundant glycerophospholipids in *E. coli*. As lipid extracts from tissues or environmental samples are much more complex, initial chemical fractionation of lipids could be used to make data analysis more tractable. Isotope ratios should be little affected by chemical separation and thus D_2_O-labeling lipidomics can be optimized to a specific ecosystem. The readout that D_2_O-labeling lipidomics enables can be applied to the study of microbial growth rates in clinical samples. It can, for example, also be applied to differentiate biologically active from inactive biomass, necromass, and contaminants.

Currently, differences in the lipid composition between microbes are already used to identify strains by chemotaxonomy ^[45,46]^. By combining large-scale lipid detection with the quantification of isotopic labeling, new applications might become possible. These include identifying microbial adaptations to drugs, determining instantaneous microbial growth rates and forecasting composition of microbial community composition after exposure to a stressor. Recording isotope labeling dynamics of lipids can help to rationalize microbial lipid function and metabolism. These efforts will benefit from related lines of research in environmental microbiology and in human physiology that measure the synthesis and turnover of lipids with isotope labeling lipidomics, mass isotopomer distribution analysis or biomarkers analysis ^[8,21,47]^.

## MATERIALS AND METHODS

### Deuterium-enriched growth medium

M9 minimal medium was prepared with 3.8 μM thiamine pyrophosphate and glucose (22.2 mM) or sodium acetate (15 mM) ^[48]^. All media were sterilized by filtration (0.2 μm). D content was adjusted by isotope dilution (measured by weight) of D_2_O (D, 99.9 at%; Cambridge Isotope Laboratories) with natural abundance water (MilliQ, EMD Millipore) of known isotopic composition. ^2^*F*_WATER_ of culture medium was measured on a DLT-100 liquid water isotope analyzer (Los Gatos Research). Samples were analyzed in three technical replicates, each comprising 10-12 injections. Samples with D abundances close to natural abundance were calibrated against standards ranging from 0.0136 % to 0.0200 % ^2^*F*_WATER_ (corresponding to δD values from −124 to +287 ‰). These were in turn calibrated against the VSMOW, GISP, and SLAP international standards ^[49]^. More enriched samples were measured against working standards made in-house, ranging from 0.050 % to 0.150 % ^2^*F*_WATER_. The presence of doubly-substituted species (D-O-D) was not taken into consideration due to fast equilibration of water molecules below 0.0150 %. Samples beyond this scale were no longer in the linear response range of the instrument, and we calculated ^2^*F*_WATER_ based on the gravimetric preparation of the medium. Their ^2^*F* values were confirmed by water isotope analysis after dilution with natural abundance water of known isotopic composition.

### *E. coli* cultures

*Escherichia coli* K-12 (FRAG1) was streaked on LB agar plates for single colonies and used to inoculate 6 mL precultures of M9 minimal medium with glucose as carbon source ^[50]^. All cultures were incubated at 37 °C and shaking at 250 rpm (Innova 44 shaker, New Brunswick Scientific). Cultures were checked by phase contrast microscopy (Axio Scope.A1, Zeiss). Optical density (OD) was measured at 600 nm wavelength (DU 800 spectrophotometer; Beckman Coulter).

To investigate the detection limits of ^2^*F*_LIPID_ and the comparison of LC-MS with GC/P/IRMS, precultures were grown for 20 hours at natural D abundance. 75 μL were used to inoculate 150 mL medium in 1 L Erlenmeyer flasks. For analysis with GC/P/IRMS the medium had a ^2^*F*_WATER_ in the range from 0.0142 % to 0.0202 % (δD −90 to +300 ‰). For analysis with LC-MS the medium had a D content of 0.0142 % to 4 % ^2^*F*_WATER_. Cultures were harvested at OD_600_ 0.2-0.3 by chilling 50 mL culture in ice and centrifugation at 4 °C for 20 min at 5,000 × g. Cell pellets were frozen in liquid nitrogen and stored at −20 °C.

For monitoring change of ^2^*F*_LIPID_ during stationary phase exit, two precultures (0.0142 % and 4 % ^2^*F*_WATER_) were centrifuged at 15 °C for 10 minutes at 15,000 × g. The pellets were resuspended in prewarmed medium (0.0142 % or 4 % ^2^*F*_WATER_) and used to inoculate 200 mL M9 glucose medium at an initial OD_600_ of 0.1. This yielded four combinations (*uU, uL, lU, lL*) of unlabeled/labeled inoculum (*u,l*) in unlabeled/labeled growth medium (*U, L*). Aliquots were incubated and sampled as described above (20 minutes: 40 mL, 50 minutes: 30 mL, 100 minutes: 30 mL and 150 minutes: 30 mL). At each time point, OD_600_ was recorded and protein content of the bacterial culture was measured via BCA protein assay (Thermo Scientific) from a cell pellet (2 mL aliquot, centrifuged at 4 °C for 2 minutes at 16,900 × g). Maximum OD_600_ was 0.8 for cultures *uU, uL* and 1.2-1.3 for cultures *lU, lL.* Before the BCA assay cells were chemically lysed (BugBuster, EMD Chemicals). Absorbance was recorded at 562 nm on a plate reader (Synergy 4, BioTek).

### Liquid chromatography mass spectrometry (LC-MS)

For LC-MS analysis lipids were extracted based on the procedure by Matyash *et al.*^*[51]*^. Cell pellets were resuspended in 0.1 % ammonium acetate to a protein concentration of 200 mg/mL (20 OD_600_ /mL). 100 μL of this suspension were added to 1.5 mL methanol, then 5 mL methyl t-butyl ether (MTBE) and a mix of standards containing PE(17:0/17:0), PG(17:0/17:0) and 14:1(3)-15:1 cardiolipin was added (Avanti Polar Lipids). After incubation in a ultrasonic bath for 1 hour, lipids (in top phase) were extracted by adding 1.25 mL water and re-extracted by addition of 2 mL MTBE/methanol/water (10:3:2.5). Samples were dried under N_2_, stored at −20 °C and dissolved in 1 mL methanol/dichloromethane (9:1) for analysis by LC-MS.

LC-MS data were collected on an Acquity I-Class UPLC coupled to a Xevo G2-S TOF mass spectrometer (Waters). Intact polar lipids were separated on an Acquity UPLC CSH C18 column (2.1 mm × 100 mm, 1.7 μm; Waters) at 55 °C following a protocol established by Waters Corporation and adapted in our laboratory ^[28]^. Samples were run in three randomized instrument replicates (injection volume 5 μL). LC-TOF-MS^E^ data was collected in positive and negative mode using electrospray ionization (ESI) with a desolvation temperature of 550 °C and source temperature of 120 °C.

Lipids were identified by the mass to charge ratio (*m/z*) of their molecular ion, their fragmentation products in positive and negative mode, and comparison to representative standards. Note that that the assignment of *sn-*1 and *sn-*2 fatty acyl positions is tentative ^[30]^. Lipids are named based on LIPIDMAPS classification ^[52]^. Table S1 provides a summary of ions and retention times used for quantification. The monoisotopic intensity alone yields incorrect values for absolute lipid concentrations in labeling experiments, where some of the signal has shifted to higher masses. So we quantified the absolute concentrations of labeled lipids by comparing the sum of all intensities of its isotopologues with those of an internal standard (Figure S6). Peaks in extracted ion chromatograms were integrated using the software MAVEN ^[53]^. Subsequent analysis was done in R ^[54]^. Models of isotopic distribution patterns were calculated using the R package Isopat (Martin Loos, EAWAG, Switzerland).

In order to calculate the generation time we fit our data to an exponential growth model. We consider that the number of lipids increases linearly with the number of cells. The number of lipid molecules at time t are given by

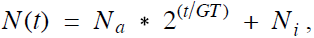

where N_a_ is the initial number of exponentially reproduced molecules, GT is the generation time and N_i_ the number of ‘inactive’ molecules at time 0. Time course data of the fractional abundance of D in the lipids (^2^*F*_LIPID_) was fitted using the equation

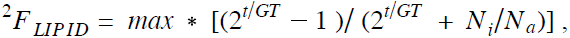

where *max* is the saturation level of labeling.

For direct infusions a stock solution of PE(18:0/18:1) in chloroform (10 g/L; Avanti Polar Lipids) was diluted to 1 μmol/L in 4:1 acetonitrile/water containing 10 mM ammonium formate and 0.1 % formic acid. H_2_ O (0.015 at% D) or D_2_O (99.9 at% D) was used for the water portion. PE(18:0/18:1) was initially dissolved in acetonitrile, later ammonium formate dissolved in water, and 0.1 % formic acid was added. Direct infusion data was collected for 4 minutes at a flow rate of 0.4 mL/min. ESI and detector settings were as in LC-MS lipidomics experiments. First the H_2_ O sample was measured, followed by washing the pumps and measuring the D_2_O sample.

Orbitrap measurements were performed at Thermo Fisher Scientific (San Jose, CA) using a Q Exactive Plus mass spectrometer operating at a full MS scan mode at a resolution of 35,000 (FWHM at *m/z* 200). The *m/z* range was set to *m/z* 150 to 2000 for negative ion mode and *m/z* 150 to 1200 for positive ion mode. The automatic gain control target value was set at 10^6^ and the maximum injection time was set at 50 ms. Chromatography was performed on a Vanquish UHPLC (Thermo Fisher Scientific), but otherwise unchanged.

### CF sputum and *S. aureus*

Sputum collection was approved by the Institutional Review Board at Children’s Hospital Los Angeles (IRB# CCI-13-00211). All patients were recruited from Children’s Hospital Los Angeles and informed consent or assent was obtained from all study participants or from a parent or legal guardian. Information about microbial community composition and fatty acid analysis of sputum samples is published elsewhere ^[42]^. Samples used for LC-MS were from Patients 1 (2nd hospitalization, day 5; method development), Patient 2 (day 2; growth rates). *S. aureus* negative controls: Patient 5 (day 1), Patient 7 (day 18 and 19).

Lipid extracts from *S. aureus* (grown in LB medium) were prepared and analyzed by LC-MS as for *E. coli*. For sputum, 10 mg of lyophilized material was extracted by the same method. The lipid extract was dissolved in 100 μL methanol and 0.2 μL were injected for routine lipidomic profiling. Detection of *anteiso-*containing phospholipids in sputum was performed by injecting 5 μL lipid extract and restricting flow into the ESI source to a retention time window of 5-8 min.

### Calculation of deuterium content in lipids (^2^*F*_LIPID_)

The mass spectra of a labeled and unlabeled lipid were compared to determine the fractional abundance of D (^2^*F*_LIPID_). We calculated the molecular weights of the two isotopologue distributions and dividing their difference by the number of C-bound hydrogen atoms. Note that we did not calculate the molecular weights using accurate masses. At 30,000 mass resolving power each signal M1, M2, etc contains isotopologues which differ slightly in mass. Measured masses have additional experimental uncertainties. For simplicity, we rather used the fact that each isotopologue must have gained a certain number of neutrons, for example M8 has gained a total of 8 neutrons from ^13^C, ^2^H etc. We used the isotopologue distribution to calculate by how many neutrons the distribution had shifted with respect to the monoisotopic mass M0. This approach eliminates inaccuracies. It also avoids the complicating fact that the mass difference between a D and ^1^H is not exactly 1.

### GC pyrolysis isotope-ratio mass spectrometry

20 mg of frozen and lyophilized cell pellet was transesterified and extracted in hexane/anhydrous methanol/acetyl chloride at 100 °C for 10 minutes ^[55]^. The extract was concentrated under N_2_. Fatty acid methyl esters (FAMEs) were first analyzed by gas chromatography mass spectrometry (GC-MS) on a Thermo-Scientific Trace / DSQ with a ZB-5ms column (30 m × 0.25 mm, film thickness 0.25 μm) and PTV injector operated in splitless mode. Peaks were identified by comparison of mass spectra and retention times to authentic standards and library data.

The δD of FAMEs was measured by gas chromatography pyrolysis isotope-ratio mass spectrometry (GC/P/IRMS) on a Thermo-Scientific DELTA^plus^XP with methane of known isotopic composition as the calibration standard ^[37]^. Chromatographic conditions were identical as for regular GC-MS, and peaks were identified by retention order and relative height. Samples were analyzed in triplicate. All data were corrected for methyl H originating from methanol by analyzing the dimethyl derivative of a phthalic acid standard, for which the δD value of ring H is known. For comparison with LC-MS, δD values were converted into fractional abundances (^2^*F*).

## ACKNOWLEDGEMENTS

We thank all reviewers for comments. We are grateful to Drs. Fenfang Wu, Reto Wijker and Jesse Allen for technical advice on the use of instrumentation, and to Dr. Ajay Kasai for CF sputum collection. LC-MS data was collected at the Caltech Environmental Analysis Center (Pasadena) and with assistance of Dr. Anastasia Kalli at Thermo Fisher Scientific (San Jose, CA). This work was made possible in parts by grants from NASA (NNX12AD93G), the National Science Foundation (1224158), the National Institutes of Health (R01HL117328). IRB was funded by CEMI (Caltech) and by the Leverhulme Trust.

## SUPPLEMENTAL INFORMATION

### Supplemental Figures S1-S6

#### Supplemental Table S1: Information about quantified lipids. (see separate file)

**Figure S1:**
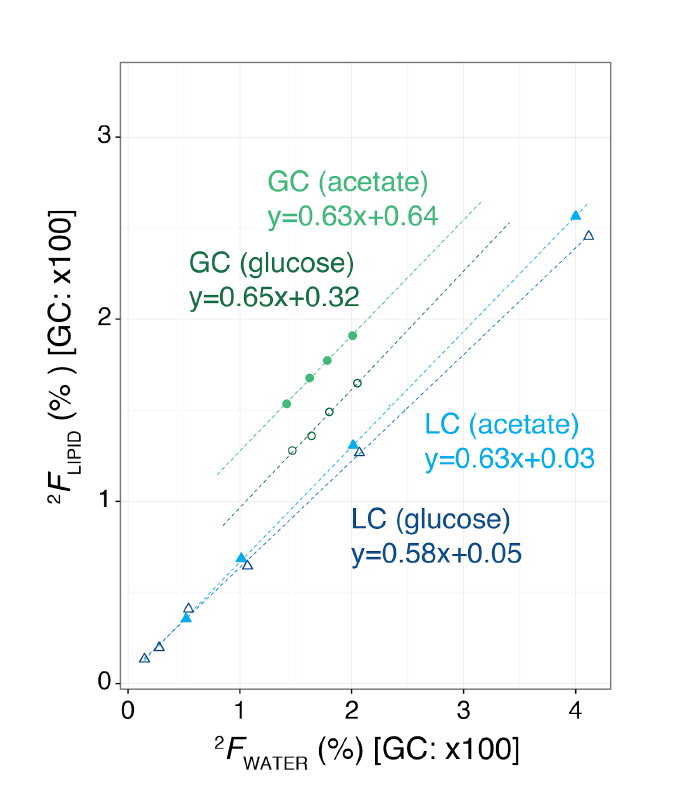
Isotope-ratio GC and LC-MS have comparable water fractionation factors. *E. coli* cultures were grown on glucose (darker color) or acetate (lighter color) as carbon sources using varying amounts of deuterium in the growth medium (^2^*F*_WATER_). For both methods deuterium content in lipids (LC) or fatty acids (GC) increases linearly with ^2^*F*_WATER_. Slopes are comparable between LC (blue triangles) and GC/P/IRMS (green circles). Data for fatty acid 16:0 (GC-MS) and PG(16:0/16:0) (LC-MS) are shown. Values for GC-MS are multiplied by a factor of 100 for easy comparison with values from LC-MS. Filled points represent glucose, empty points acetate cultures.

**Figure S2:**
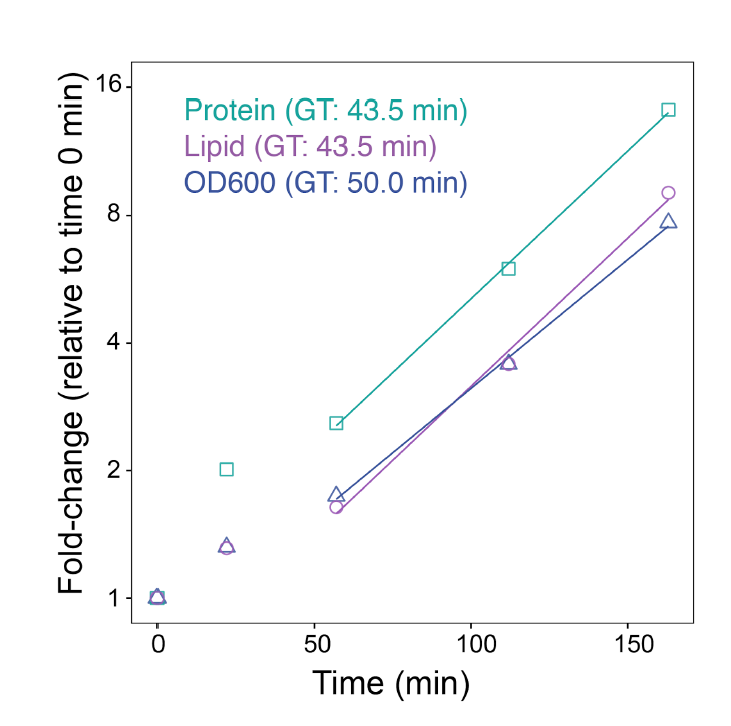
Estimates of generation time (GT) from lipid, protein and OD_600_measurements are similar. Protein abundance grew disproportionately during initial cell expansion, likely owing to the fact that stationary phase cells were smaller than those in exponential phase. In exponential phase, the three methods yield generation times of 43.5 minutes for protein (green squares) as well as lipids (magenta circles) and 50 minutes for OD_600_(blue triangles). Data from a single culture (*uU* culture) are shown.

**Figure S3:**
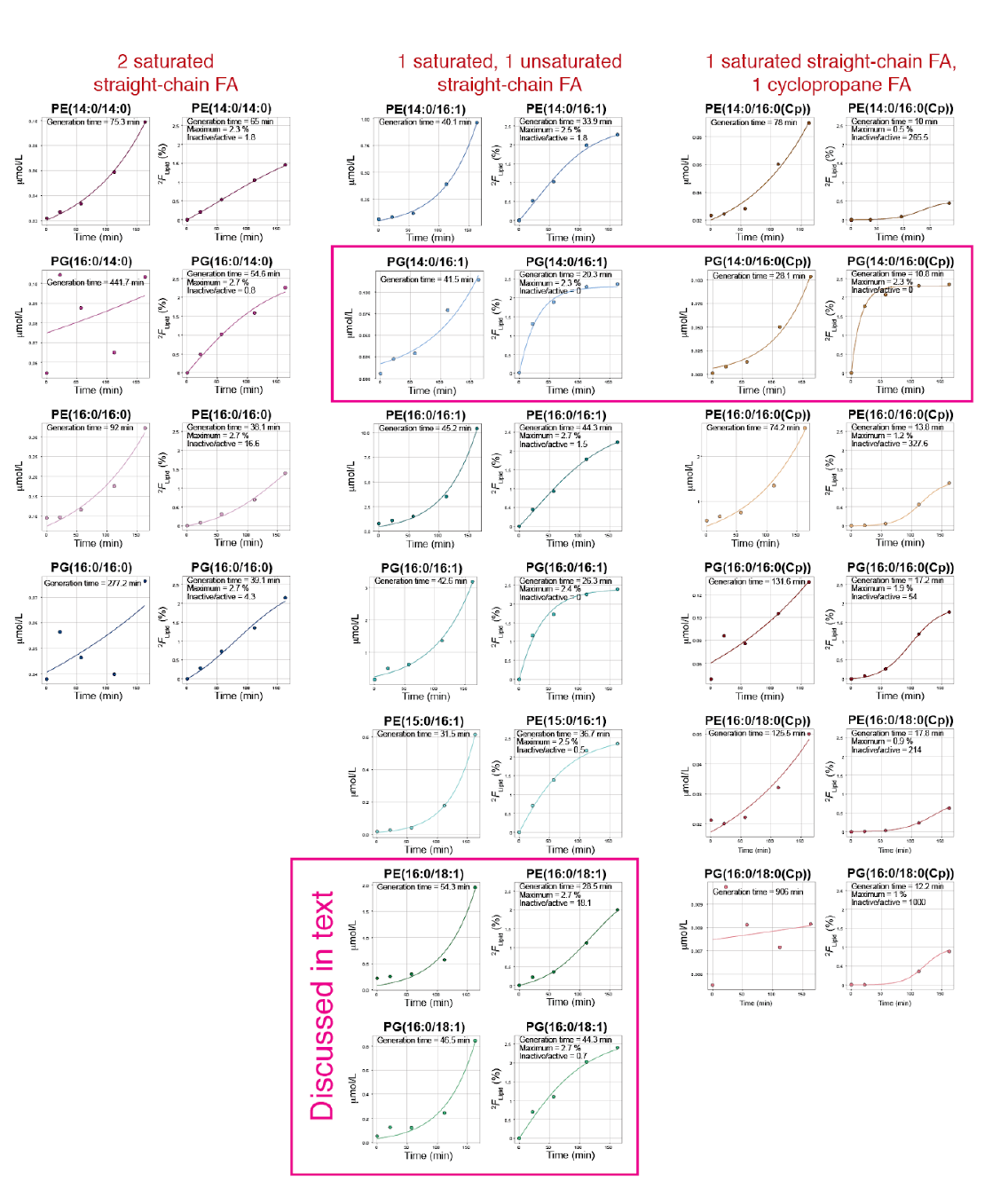
Overview of *E. coli* PE and PG lipid production during the transition from stationary phase into exponential phase. Phospholipids from culture *uL* were quantified by the summed signals of their isotopologues and compared to their deuterium abundance (^2^*F*_LIPID_). Lipids were grouped based on their fatty acyls, because this correlated with labeling dynamics. Notable exceptions of common trends were observed for example with PE(16:0/18:1) and PG(14:0/16:0(Cp)).

**Figure S4:**
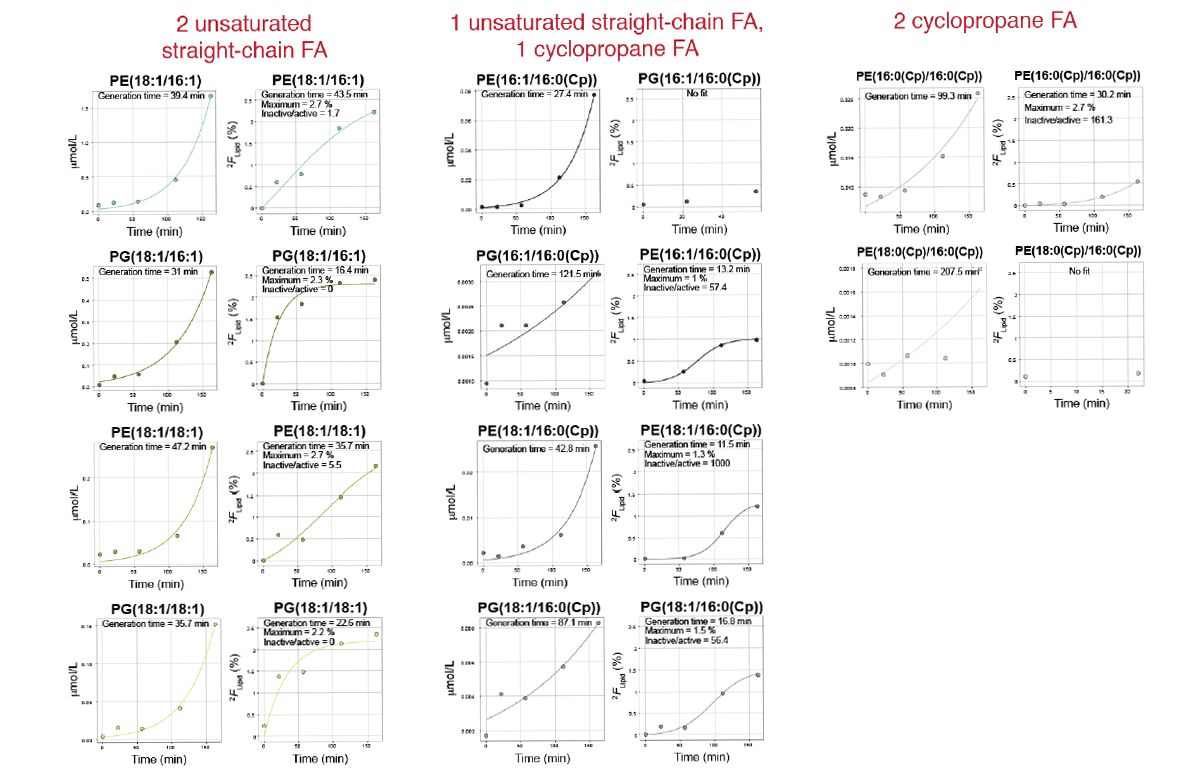
Overview of *E. coli* PE and PG lipid production during the transition from stationary phase into exponential phase (continued).

**Figure S5:**
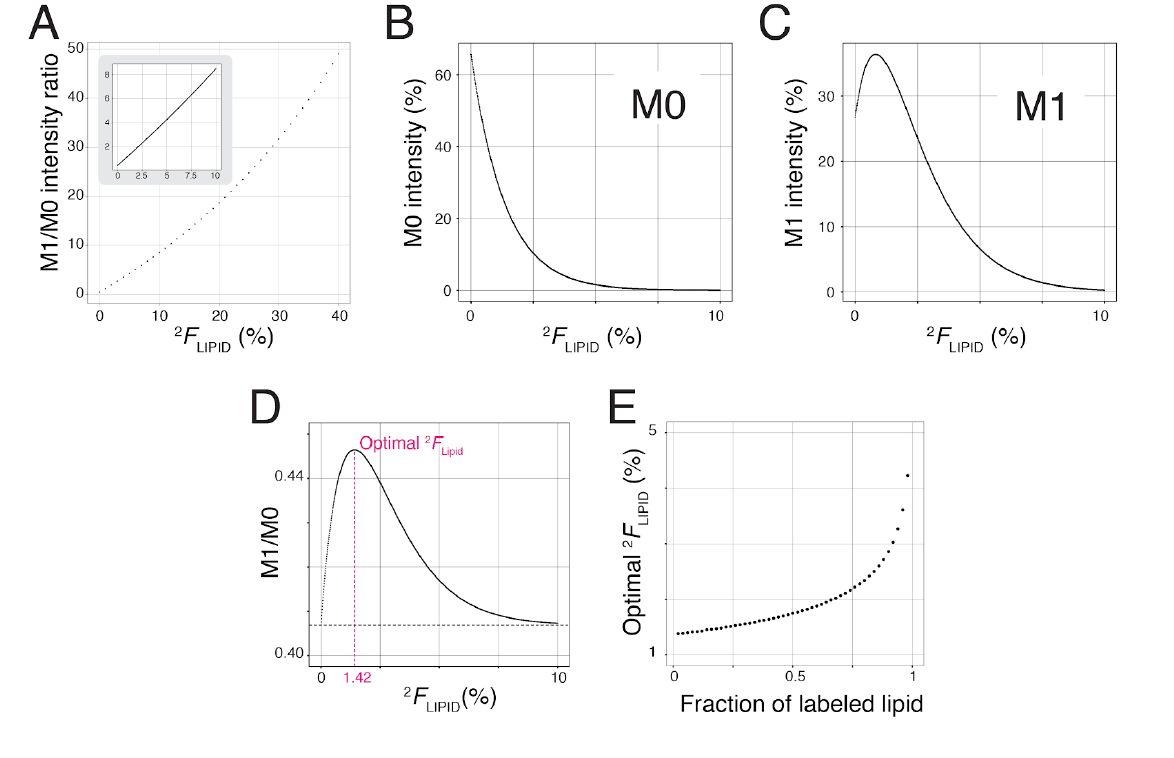
Deuterium content in lipids can be quantified using the ratio of M1 and M0 intensities. The sum formula of the phospholipid PE(16:0/16:1), C_37_H_73_NO_8_P, was used to model the behaviour of M1 and M0 isotopologues. **(A)** Deuterium abundance in lipid (^2^*F*_LIPID_) was altered from 0 to 40 at%. M1/M0 rises approximately linearly at ^2^*F*_LIPID_< 10 at% (inset). **(B and C)** Theoretical modeling of the proportion of M0 and M1 intensity compared to all isotopologues with varying ^2^*F*_LIPID_. **(D)** M1/M0 does not change linearly with increasing ^2^*F*_LIPID_of labeled lipid when unlabeled material is also present. The calculation assumed a mixing of 90 mol% unlabeled lipid and 10 mol% deuterium labeled lipid of varying deuterium abundance ^2^*F*_LIPID_(plotted on x-axis). In this scenario an optimum is observed when the labeled lipid portion has a ^2^*F*_LIPID_value of 1.42 %. **(E)** This optimal ^2^*F*_LIPID_increases the more labeled lipid is produced. When 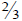 labeled lipid and 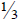 unlabeled lipid is assumed, the optimal ^2^*F*_LIPID_is about 2%. The optimum also depends on the number of hydrogens in a lipid.

**Figure S6:**
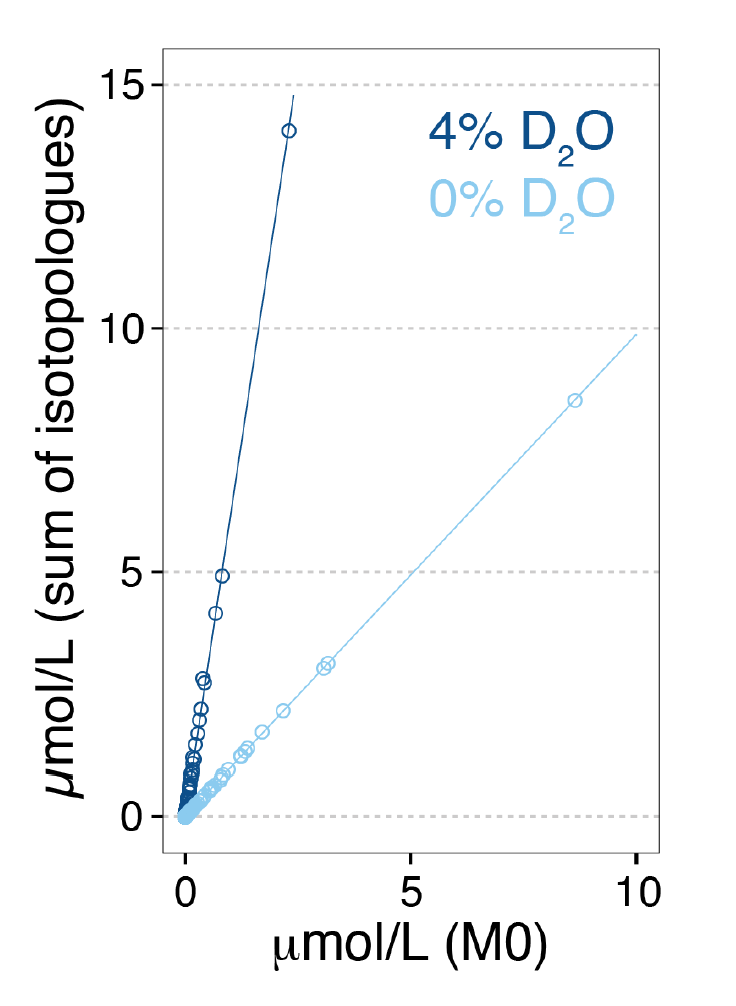
Absolute abundance of labeled lipids can quantified with the summed intensities of isotopologues. PE and PG lipids of two *E. coli* cultures unlabeled or full labeled in 4 % D_2_O medium, were quantified based on their monoisotopic signal or the sum of all isotopologues. A linear relationship between these two quantification methods were observed for lipids from both cultures. This suggests that, in experiments with little ion suppression, D-labeled lipids can be accurately quantified by comparing the sum of isotopologue intensities of lipids and internal standards. In the unlabeled culture the slope was 0.977±0.034 (1 σ) for PE, 0.980±0.037 for PG and 1.05±0.118 for CL.

